# Heterozygous missense *RAD21* variant in a peripheral sclerocornea pedigree

**DOI:** 10.1101/547547

**Authors:** Bi Ning Zhang, Tommy Chung Yan Chan, Pancy Oi Sin Tam, Yu Liu, Chi Pui Pang, Vishal Jhanji, Li Jia Chen, Wai Kit Chu

## Abstract

**Background:** Sclerocornea is a rare congenital disorder characterized with cornea opacification. We identified a heterozygous missense *RAD21* variant in a non-cons anguineous Chinese family with multiple peripheral sclerocornea patients spanning across three generations inherited in an autosomal dominant manner.

**Methods:** Comprehensive ophthalmic examinations were conducted on all 14 members. Whole exome sequencing was used to identify the genetic alterations in the affected pedigree members. Lymphoblastoid cell lines (LCLs) were established using blood samples from all members. Cleavage of RAD21 protein was quantified in these cell lines.

**Results:** All affected individuals showed features of scleralization over the peripheral cornea of both eyes. Mean horizontal and vertical corneal diameter were significantly decreased in the affected members. Significant differences were also observed on mean apex pachymetry between affected and unaffected subjects. A *RAD21*^*C1348T*^ variant was co-segregated with affected members. Both the wild-type allele and the missense variant were expressed at the mRNA level. This variant caused RAD21 R450C substitution at the separase cleavage site, which led to reduced RAD21 cleavage.

*Conclusion:* We believe this is the first report of genetic variant in sclerocornea without other syndromes. Further work is needed to confirm the RAD21^R450C^ variant with sclerocomea.

## Introduction

Sclerocornea is a rare corneal opacification disease. The transition between the sclera and cornea is indistinct in sclerocornea cases (1, 2). If the opaque area is too larger to affect vision, corneal transplantation is the only way to restore vision in sclerocornea patients. Given the shortage of cornea for transplantation and the high failure rate of keratoplasty for young children (3), better understanding of the disease mechanism of sclerocornea is needed so that alternative treatments can be developed.

Cohesin is comprised of two structural maintenance of chromosomes subunits, SMC1 and SMC3, and a kleisin subunit RAD21 to form a ring-structure protein complex that encircles sister chromatids and prevents premature sister chromatid separation (4, 5). At centromeres and telomeres, before the onset of anaphase, the active separase cleaves RAD21 proteolytically after the arginine at positions 172 (Arg172) and 450 (Arg450) (6). These cleavages on RAD21 open the cohesin complex to allow sister chromatid separation. Rad21 has been found to be highly expressed in both postnatal developing cornea and adult cornea in mice compared with other ocular tissues, suggesting potential roles of Rad21 in cornea development and cornea diseases (7).

In this study, we identified a Chinese family with multiple peripheral sclerocornea patients spanning three generations. We discovered a heterozygous variant at one of the separase cleavage sites of RAD21, R450C, associated with peripheral sclerocornea. And this variant led to reduced separase cleavage of RAD21.

## Materials and methods

### Patient consent

This study adhered to the tenets of the Declaration of Helsinki and was approved by the institutional review board of Kowloon Central Cluster, Hospital Authority, Hong Kong (KC/KE-15-0223/ER-1). Consent forms were signed by participants from this peripheral sclerocornea pedigree.

### Clinical assessments

Visual acuity assessment, refraction, slit-lamp examination, intraocular pressure measurement by Goldmann applanation tonometer and optic disc assessment were performed (Table 1). Axial lengths were determined with IOL Master V.5 (Carl Zeiss Meditec, USA). Corneal topography, pachymetry, anterior chamber depth, and anterior chamber angle were assessed using a Casia SS-1000 swept source OCT (Tomey, Japan). Retinal nerve fiber layer was evaluated with a Cirrus HD-OCT spectral domain OCT (Carl Zeiss Meditec). Fasting serum lipid profile was performed for all 14 members who attended the clinic.

**Table 1.**
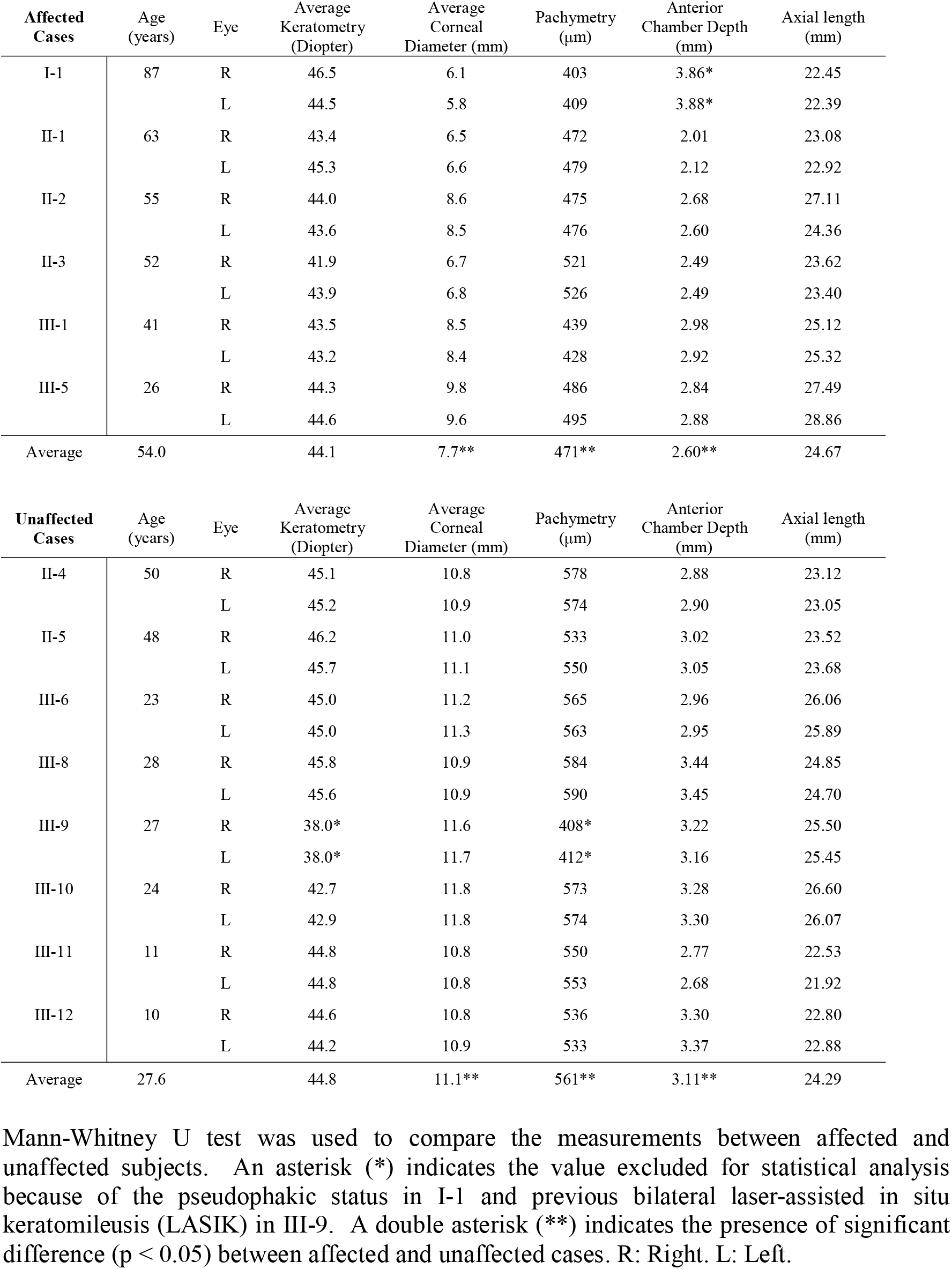
Ophthalmic examinations of a Chinese family with peripheral sclerocornea.

### Whole-exome sequencing (WES)

WES analysis (Macrogen) was performed in four affected cases and two unaffected cases. Genomic DNA was extracted using QIAamp DNA Blood Mini Kit (Qiagen, Germany) and exome DNA was enriched with an Agilent SureSelect Human All Exon V5 kit (Agilent Technologies, USA). Paired-end sequencing was performed on Illumina HiSeq 4000 (Supplementary Table 1 and 2).

To identify functional variants that were associated with sclerocornea, raw sequencing reads were mapped onto human reference genome (hg19) using Burrows-Wheeler Aligner (BWA) (8) (Supplementary Table 3). Both SAMtools (9) and Genome Analysis Toolkit (GATK) (10) were used to call SNPs and indels. Around 77,000 variants were obtained for each case. Among all the variants, 28,593 variants that appeared in all the four affected members were filtered using an in-house pipeline to obtain 3,744 variants within protein coding regions. 2,838 variants were excluded since two unaffected members also carried them. Remaining variants were further filtered using the autosomal dominant disease model, the dbSNP135 and the 1000 Genomes datasets to exclude population associated variants. Since sclerocornea is a rare disease, a minor allele frequency threshold at 0.0001 was applied to obtain 15 rare variants. Published data were used to further examine these 15 variants (11). 5 non-synonymous SNPs were excluded since they appeared in a Han Chinese control group of 190 unrelated controls who had received comprehensive ophthalmological examinations and showed no features of sclerocornea. 15 candidate variants were obtained and summarized in Supplementary Table 1.

### Lymphoblastoid cell lines (LCLs)

Fresh blood was collected and lymphocytes were isolated using Ficoll-Paque (GE Healthcare, USA). Lymphocytes were transformed with B95-8 Epstein-Barr virus (ATCC, ATCC) according to the described protocol (12). 2μg/mL cyclosporin A (Sigma-Aldrich, USA) was added to the culture medium until cells formed a rosette morphology. LCLs were cultured with Iscove’s Modified Dulbecco’s Medium supplemented with 20% fetal bovine serum, 100U/mL penicillin and 100μg/mL streptomycin (Gibco, USA) at 37 °C in a 95% air-5% CO2 incubator.

### LCLs characterization

Allele specific gene expression analysis was conducted with data from RNA sequencing (Macrogen, South Korea). Western blot was conducted with anti-RAD21 antibody (ab42522, Abcam, UK) and goat anti-rabbit antibody (sc-2004, Santa Cruz, USA). GAPDH (G9295, Sigma-Aldrich) was used as a loading control.

## Results

### A Chinese peripheral sclerocornea pedigree spanning three generations

A non-consanguineous Chinese family with peripheral sclerocornea was identified from the Hong Kong Eye Hospital. 8 cases (6 males and 2 females) of peripheral sclerocornea spanning across 3 generations were found in this family (Figure 1A). Fourteen members were examined with slit-lamp and 6 cases (I-1, II-1, II-2, II-3, III-1 and III-5) were identified to have peripheral sclerocornea (Figure 1B). Two additional cases in the third generation were identified to have peripheral sclerocornea solely based on histories. All affected subjects had bilateral sclerocornea. The degree of peripheral scleralization was similar in both eyes, despite the variations that existed among individuals. Peripheral sclerocornea was not noted in the examined unaffected subjects (II-4, II-5, III-6, III-8, III-9, III-10, III-11 and III-12). For all 14 members, the overall mean±standard deviation and median age was 38.9±21.6 years (range: 10-87 years) and 34.5 years respectively. Comparing the affected and unaffected members, the mean age was 54.0±20.6 years (range: 26-87 years; median 53.5 years) and 27.6±14.8 years (range: 10-50 years; median 25.5 years) respectively (p=0.020). In the affected subjects, two members (II-1 and II-3) had cataract and one member (I-1) had age-related macular degeneration.

**Figure 1.**
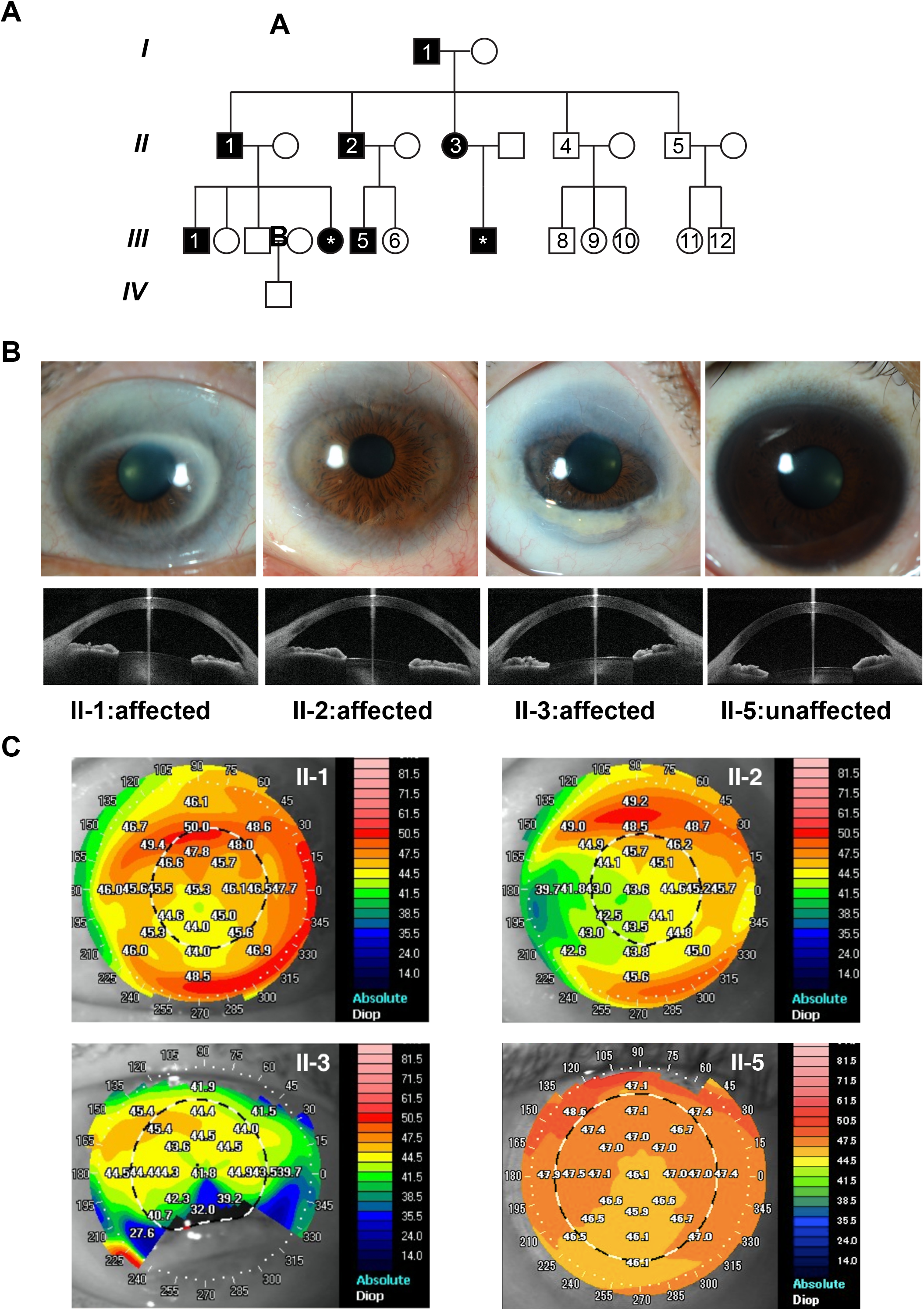
**A.** The pedigree of a family showing autosomal dominant inherited peripheral sclerocornea. Black and white indicate affected and unaffected members respectively, whereas squares and circles represent male and female respectively. Roman numbers indicate the generations and Arabic numbers show the family members who gave consent to clinical exams and genetic analysis. **B.** Photographs of the eyes from three affected members (II-1, II-2, II-3) and one unaffected individual (II-5). In the eyes of affected members, white regions were observed on the periphery of corneas, indicating the peripheral sclerocornea. These white regions were not observed in the unaffected eye. Anterior segment imaging with the swept source anterior segment OCT in selected family members (II-1, II-2, II-3, and II-5). Only horizontal section images of each subject are shown. No abnormality of the anterior chamber angle is found in affected and unaffected individuals. **C.** Corneal topography of selected family members. Keratometry readings are shown on the corneas and represented by color codes as shown on the right. Top left: the image for the left eye image of II-1 (affected subject); Top right: The left eye image of II-2 (affected subject); Bottom left: The image for the right eye image of II-3 (affected subject); Bottom right: Right eye image of II-5 (unaffected subject).

### Corneal diameter was influenced by peripheral sclerocornea

11 of the 14 members had best-corrected visual acuity (BCVA) of ≥20/30 in either eye. Three affected members had BCVA of < 20/30. The mean logMAR BCVA was 0.17±0.22 (range: 0–0.7) for all the 14 members of the family. Only myopia or mild hypermetropia was noted. The spherical equivalent ranged from -15.0D to +2.5D for all the 14 members. Comparing the affected and unaffected subjects, the mean spherical equivalent was -4.70±5.97 diopters for the 6 affected members and -1.57±2.89 diopters for the 7 unaffected members (p=0.138). The mean horizontal and vertical corneal diameter for the affected subjects was 8.7±1.4mm and 6.6±1.7mm, respectively. The corresponding diameter was 11.6±0.4mm and 10.6±0.5mm for the unaffected subjects. Significant differences in both horizontal and vertical corneal diameter were observed between affected and unaffected family members (p=0.013 and p=0.001 for horizontal and vertical diameter, respectively).

In all the 14 members, there were no corneal endothelial guttate, deep central corneal stromal opacities, iris abnormalities, iridocorneal adhesion or shallow anterior chamber. Anterior chamber and angle analysis did not reveal any abnormality (Figure 1B). An open angle was identified in all cases. The intraocular pressure of both affected and unaffected subjects was comparable; the corresponding mean value was 14.6±1.1mmHg (range, 12 to 18mmHg) and 16.3±1.3mmHg (range, 14 to 19mmHg) (p=0.073) respectively. There were no features of glaucoma either on optic disc assessment or retinal nerve fiber layer evaluation.

### Mean apex pachymetry differs significantly between the affected and unaffected members

The keratometric readings of the 6 affected subjects ranged from 41.9 to 46.5 diopters (mean, 44.1±0.9 diopters), while the readings of the unaffected 7 subjects ranged from 42.7 to 46.2 diopters (mean, 44.8±1.0 diopters) (Figure 1C) (p=0.153). III-9 was excluded in this analysis because of the previous bilateral laser-assisted in situ keratomileusis (LASIK). The keratometry readings of affected subjects were within normal range and were not different from the unaffected subjects of the same family (Table 1). The mean preoperative keratometry for the affected subject who underwent bilateral cataract surgery (I-1) was 46.5D for right eye and 44.5D for left eye whereas the postoperative mean keratometry values were 47.3D for right eye and 45.2D for left eye. The mean apex pachymetry of the 6 affected members was 471±39μm and the corresponding value of the unaffected members was 561±19μm, which differed significantly (p=0.003). The mean anterior chamber depths were 2.60±0.60mm and 3.11±0.25mm for the affected and unaffected members, respectively (p = 0. 013). Axial length of the affected subjects varied between 22.39 and 28.86mm (mean, 24.67±2.14mm), while the axial length of the unaffected subjects ranged from 21.92 to 26.60 mm (mean, 24.29±1.56mm) (p=0.796). None of the family members had mental deficits, metabolic syndromes or systemic abnormalities except for hypertension in 3 cases. Serum lipid profile was normal for all 14 cases. The average keratometry and axial length in the affected subjects were not particularly hypermetropic as expected in cornea plana (13).

### Co-segregation of heterozygous variant *RAD21* c.C1348T with peripheral sclerocornea in the pedigree

We performed whole exome sequencing (WES) in four affected (I-1, II-1, II-2, II-3) and two unaffected (II-4, II-5) members (Figure 1A, Supplementary Table 1, 2 and 3). By using a stepwise filtering method, 15 rare candidate variants from 10 genes were obtained (Supplementary Table 1). Among these 15 variants, the missense variant *RAD21* c.C1348T (p.R450C) showed complete co-segregation with the peripheral sclerocornea in the pedigree. Sanger sequencing results confirmed that the affected members carry both wild-type and variants (Figure 2A).

**Figure 2.**
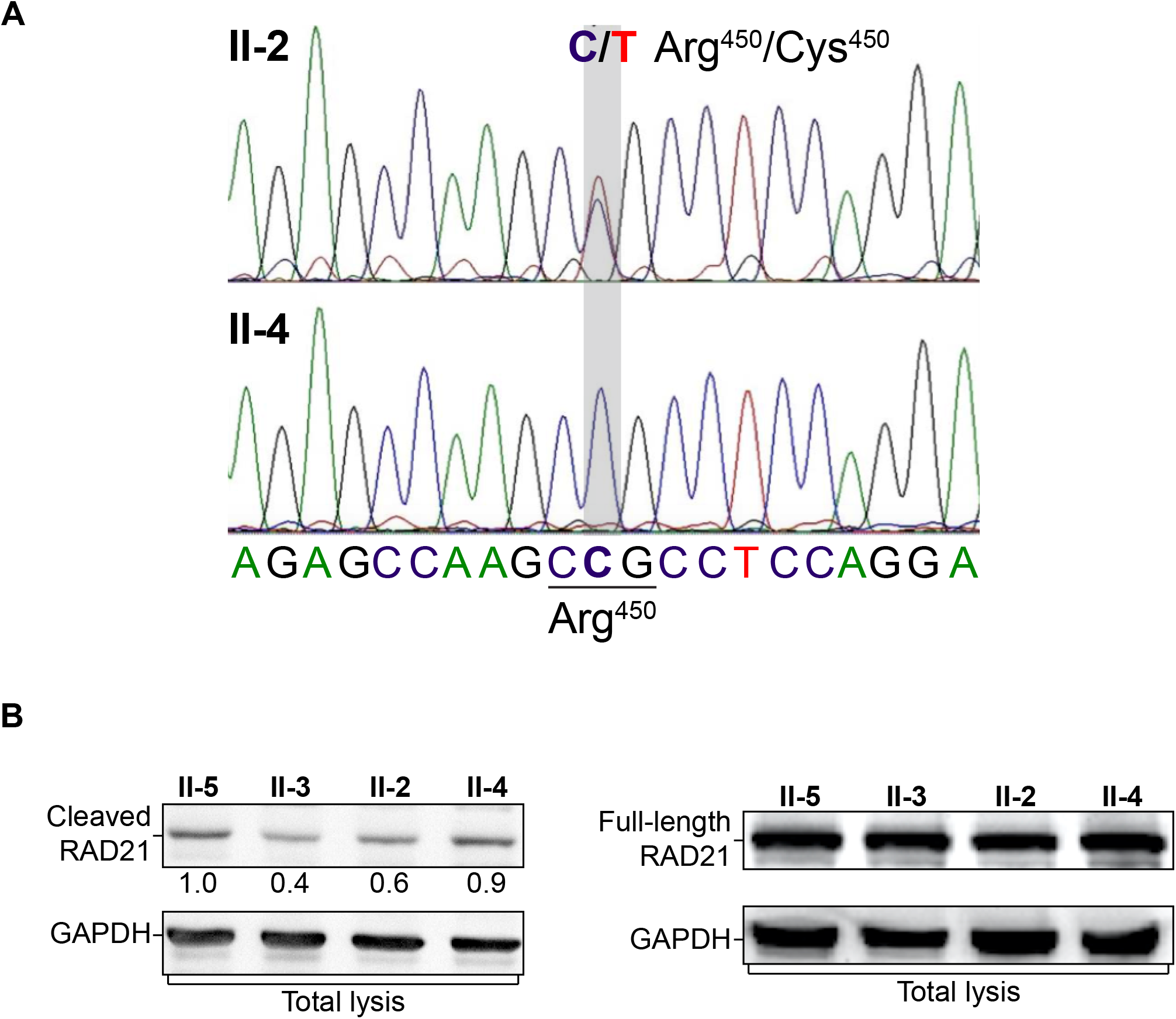
**A.** Sanger sequencing spectra of *RAD21* alleles in affected and unaffected members. In the locus, the affected sample (II-2) showed two sequencing peaks, indicating both signals of dC and dT. The unaffected sample (II-4) showed only one peak of dC. **B.** Total protein extracts were analyzed using Western blotting. The intensities of the separase cleaved RAD21 bands decreased in affected members. Normalized levels of cleaved RAD21 relative to that of II-5 are shown under the Western blot image. Protein levels of the full-length RAD21 were comparable. GAPDH is shown as loading controls.

### RAD21 variant was less prone to separase mediated cleavage

The Arg450 is one of the two separase cleavage sites on RAD21, we thus examined whether the R450C variant would affect the separase cleavage on RAD21. We found that the RAD21 c.C1348T allele counted for 55% and 52% of the total *RAD21* expression levels in the affected member II-2 and II-3, respectively, whereas unaffected LCLs only express the wild-type allele. To examine whether the heterozygous RAD21 R450C variant could influence the separase cleavage on RAD21, we used a commercial antibody to detect the 31.5kDa RAD21 fragment (amino acid 172-450) generated by separase cleavages on Arg172 and Arg450 in LCLs. Although the full-length protein levels of RAD21 show no difference between the affected and unaffected LCLs (Figure 2B), the RAD21^172-450^ protein levels were lower in the affected group than that in the unaffected group, suggesting less separase cleavage on the RAD21 C450 site (Figure 2B).

## Discussion

In this study, we performed systematic ophthalmic exams on 14 family members of this pedigree. The results showed that all affected members have non-transparent corneal rim and other features of peripheral sclerocornea. These clinical results indicated that peripheral sclerocornea of this pedigree is not complicated with other syndromes, thus allowing us to identify molecular causes for peripheral sclerocornea.

Although a few familial hereditary sclerocornea were reported (14, 15), sclerocornea-associated gene mutations have only been reported in individual sclerocornea patients. For example, heterozygous mutations in *RAX* were found in a single unilateral anophthalmia and sclerocornea case (16). Mutations in *FOXE3* were reported to cause anterior segment ocular dysgenesis, including sclerocornea, microphthalmia, aphakia and absence of iris (17, 18). Mutations at the chromosomal locus Xp22.3 were associated with a X-chromosome linked sclerocornea, microphthalmia and dermal aplasia syndrome (19). Although these mutations were reported associated with sclerocornea, clinical outcomes of these mutations are complicated with other syndromes. In addition to rs1051321465 (R450H), RAD21 R450C has been included as rs130282588 in the dbSNP database. However, neither population nor clinical association was reported with these two SNPs.

RAD21 contains two reported separase cleavage sites and mutations of these two sites to alanine would lead to sister chromatids mis-segregation, cell cycle defects and aneuploidy (6). The variant identified in our pedigree is located on Arg450, one of the two separase cleavage sites and our Western blotting result revealed less separase cleavage product on the RAD21 carrying this variant, suggesting R450C affects separase cleavage. Further studies and additional patients will be useful for defining the genotype-phenotype correlations.

## Declarations

### Funding

This work was supported by The Chinese University of Hong Kong Direct Grant (Project 4054284, to W.K.C.), National Natural Science Foundation of China (Project 81770903, to W.K.C.) and the Endowment Fund for Lim Por-Yen Eye Genetics Research Centre, Hong Kong.

### Author Contributions

B.Z., T.C.Y.C., V.J., L.J.C., and W.K.C. conceived and designed this study. T.C.Y.C. and V.J. identified the pedigree and performed clinical examinations. P.O.S.T. and B.Z. performed WES analysis. B.Z. generated and characterized LCLs. B.Z., Y.L., T.C.Y.C, and W.K.C. wrote the manuscript. C.P.P., V.J., L.J.C., and W.K.C. supervised the project.

### Competing interest statement

All authors have no competing interests to declare.

