## Supplementary Tables for "Heterozygous missense *RAD21* variant in a peripheral sclerocornea pedigree"

**Supplementary Table 1** Information of the 10 candidate variants resulting from filtering of exome sequencing data for the peripheral sclerocornea pedigree.

| **No.** | **Gene Symbol** | **mRNA** | **cDNA change** | **dbSNP** | **Protein change** | **Clinical**  **Significance** | **Reported Population** |
| --- | --- | --- | --- | --- | --- | --- | --- |
| **1** | ***RAD21*** | **NM_006265** | **c.C1348T** | **rs1301282588** | **p. R450C** | **NA** | **NON** |
| 2 | *SQSTM1* | NM_001142299 | c.C11T | rs763040103 | p.S4F |  |  |
|  |  | NM_003900 | c.C263T |  | p.S88F | NA | NON |
|  |  | NM_001142298 | c.C11T |  | p.S4F |  |  |
| 3 | *RAPGEF5* | NM_012294 | c.141_143del | NA | p.47_48del | NA | NA |
| 4 | *PRICKLE4* | NM_013397 | c.863_864insTCT | rs76510495 | p.L288delinsLL | NA | NA |
| 5 | *ANGPT2* | NM_001147 | c.T704C | rs149699486 | p.V235A |  |  |
|  |  | NM_001118888 | c.T548C |  | p.V183A | NA | NON |
|  |  | NM_001118887 | c.T704C |  | p.V235A |  |  |
| 6 | *MNT* | NM_020310 | c.695_695+1insGT | rs58552657 |  | NA | NA |
| 7 | *REXO1* | NM_020695 | c.1741_1742insTCC | rs3052937 | p.S581delinsSS | NA | NA |
| 8 | *OR7G3* | NM_001001958 | c.922_923insATACC | rs3029651 | p.I308fs | NA | AF* |
| 9 | *FAM98C* | NM_174905 | c.1047_1048insAAG | rs59917662 | p.K349delinsKK | NA | NON |
| 10 | *HSPBP1* | NM_012267 | c.76_77insGGCGGCGGC | rs10701478 | p.G26delinsGGGG | NA | NON |
|  |  | NM_001130106 | c.76_77insGGCGGCGGC |  | p.G26delinsGGGG |  |  |

NA: not available in dbSNP;

NON: non-association in dbSNP;

*AF: African 0.711, East Asian 0.365, Europe 0.314, South Asian 0.34, American 0.24.

**Supplementary Table 2** Summary of original exome sequencing data of the peripheral sclerocornea pedigree.

| **Data** | **I-1** | **II-1** | **II-2** | **II-3** | **II-4** | **II-5** |
| --- | --- | --- | --- | --- | --- | --- |
| Number of raw reads (M) | 58.9 | 92.5 | 80.0 | 62.2 | 77.0 | 70.0 |
| Average read length (bp) | 101 | 101 | 101 | 101 | 101 | 101 |
| Raw data yield (Gb) | 5.9 | 9.3 | 8.1 | 6.3 | 7.8 | 7.1 |
| Number of reads mapped to the genome (M) | 58.8 | 92.2 | 79.8 | 62.0 | 76.8 | 69.8 |
| Fraction of uniquely mapped bases on target (%) | 87.6% | 87.5% | 87.7% | 87.7% | 87.8% | 87.2% |
| Data mapped to target region (M) | 34.4 | 53.5 | 46.2 | 36.2 | 45.1 | 40.4 |
| Mean depth of target region (fold) | 44.7 | 69.6 | 60.2 | 47.0 | 58.8 | 52.4 |
| Coverage of target region (%) | 95.9% | 96.1% | 96.1% | 95.8% | 96.0% | 96.1% |
| Target region with depth of more than 10 times (%) | 90.0% | 92.0% | 91.7% | 90.4% | 91.4% | 91.1% |

**Supplementary Table 3** Summary of detected variants of the peripheral sclerocornea pedigree.

| **Variants** | **I-1** | **II-1** | **II-2** | **II-3** | **II-4** | **II-5** |
| --- | --- | --- | --- | --- | --- | --- |
| Number of SNPs | 76016 | 77937 | 77883 | 76800 | 77698 | 77649 |
| Number of coding SNPs | 20840 | 20572 | 20740 | 21087 | 20834 | 20928 |
| Number of synonymous SNPs | 10812 | 10719 | 10804 | 10981 | 10804 | 10866 |
| Number of nonsynonymous SNPs | 9475 | 9325 | 9402 | 9556 | 9483 | 9506 |
| Number of Indels | 7509 | 8170 | 8032 | 7610 | 7996 | 7754 |
| Number of coding Indels | 414 | 438 | 427 | 433 | 433 | 433 |
